# Distinct Biological and Biomechanical Features in TMJ and Knee Cartilages

**DOI:** 10.64898/2026.02.22.707211

**Authors:** Jingyu He, Hong Colleen Feng, Junxi Guo, Julia Raulino Lima, Mohammadreza Vatankhah, Fangzhou Bian, Holly Katherine Johnson, Courtney Leong, Laura Voskanyan, Jantzen Blaine Ebreo, Parth Kotak, Glenn Thomas Clark, Anette Vistoso Monreal, Amy E Merrill, Jian-Fu Chen, Jian Xu, Zhaoyang Liu

**Affiliations:** Center for Craniofacial Molecular Biology, Herman Ostrow School of Dentistry, University of Southern California, Los Angeles, CA 90033, USA; Mechanical Surface Characterization Division, Anton Paar USA Inc., 2824 Columbia Street, Torrance, CA 90503; Department of Orthopaedic Surgery, Keck School of Medicine, University of Southern California, Los Angeles, CA 90033, USA; Orofacial Pain and Oral Medicine Center, Herman Ostrow School of Dentistry, University of Southern California, Los Angeles, CA 90089, USA; Department of Stem Cell Biology and Regenerative Medicine, Keck School of Medicine, University of Southern California, Los Angeles, CA 90033, USA

**Keywords:** Temporomandibular disorders (TMD), Cartilage, Extracellular matrix (ECM), Biomechanics, Stem cells, Joint disease

## Abstract

The temporomandibular joint (TMJ) and the knee joint are two of the most frequently used joints in the body, with the mandibular condylar cartilage (MCC) and the articular cartilage (AC) covering the joint bone surfaces, respectively. Compromised MCC functions lead to various temporomandibular disorders (TMD), including TMJ osteoarthritis (TMJ OA); however, the mechanisms governing MCC homeostasis and its biomechanical properties are still poorly understood. In this study, we comprehensively compared the biological and biomechanical features of the MCC and AC in mice. Histological analysis on P1, P21, 3-month, and 10-month mice revealed the most drastic structural differences between MCC and AC at occlusion establishment (P21), with MCC found to be more susceptible to age-associated cartilage degeneration. Immunostaining revealed differentially distributed cartilage extracellular matrix components in MCC and AC, including collagen type I, II, and X, and highly enriched expression of several key transcriptional factors at the posterior region of the MCC, including sex determining region Y-box 9 (SOX9), runt-related transcription factor 2 (RUNX2), and scleraxis (SCX). The posterior MCC also houses a group of long-lasting, slow-proliferative cells, as evidenced by the BrdU/EdU incorporation assay, suggesting the presence of a potential stem/progenitor cell niche at the posterior TMJ. Unbiased nanoindentation analysis revealed distinct biomechanical features between these joint cartilages. MCC exhibits a significantly lower elastic modulus (E_IT_) than AC, with the highest E_IT_ observed at the anterior TMJ, which is oppositely associated with the fibrous layer thickness, but positively correlated with the ratio of the collagen type X-positive matrix in the cartilaginous layer. Altogether, this study provides a basic understanding of the biological and biomechanical features of the cartilaginous tissues in two important joints, which may facilitate our understanding of the physiology in the TMJ and knee joint, and support the applications of mouse models to study TMJ dysfunctions.

## Introduction

The knee joint and the temporomandibular joint (TMJ) are two of the most frequently used joints in the body. Although both are diarthrodial hinge joints, they differ structurally: the knee contains medial and lateral menisci, whereas the TMJ features a fibrocartilaginous articular disc (Bielajew et al. 2021). Each joint relies on specialized cartilage adapted to distinct mechanical environments. Knee articular cartilage (AC) is a hyaline cartilage with an extracellular matrix (ECM) rich in proteoglycans and type II collagen, lining the tibiofemoral interface (Pueyo Moliner et al. 2025), while the mandibular condylar cartilage (MCC) in TMJ exhibits a bilayer structure, with a top fibrous layer overlaying a deeper cartilaginous layer (Kuroda et al. 2009). These differences reflect joint-specific biomechanical demands, as the knee joint frequently sustains weight-bearing loads, including walking, running, and jumping, whereas the TMJ resists dynamic, multidirectional forces during jaw movement, such as chewing, speaking, and breathing (Bielajew et al. 2021; Kuroda et al. 2009; Petitjean et al. 2023).

Despite structural and functional differences, both joints are susceptible to degenerative disorders. Knee osteoarthritis (OA), characterized by cartilage degradation, bone remodeling, osteophyte formation, joint inflammation, and pain, affects about 43% of adults over 40 (Tang et al. 2025). Temporomandibular disorders (TMD), an umbrella term to describe a wide variety of TMJ pathologies, affect 15–32% of adults (Bielajew et al. 2021; Tang et al. 2025). Approximately 60% of TMD patients develop TMJ OA between the ages of 20 and 40, leading to cartilage loss, jaw pain, and limited jaw function (Melou et al. 2022). Despite its high prevalence, TMJ OA remains poorly studied, largely due to TMJ’s structural complexity and our limited mechanistic insights. It is also challenging to leverage knee OA animal models for TMJ OA study because of the distinct biological and biomechanical differences between these joints.

In this study, we comprehensively compare the biological and biomechanical properties of MCC in the TMJ and AC in the knee joint in mice by assessing histological features, ECM composition, expression pattern of key transcriptional factors for cartilage homeostasis, retention pattern of the slow-cycling/progenitor-like cells, and biomechanical properties, to facilitate our understanding of the similarities and differences between the TMJ and knee joint, with a specific effort to highlight the unique structure-function relationship in the MCC. Our findings support the concept that the bilayer structure of MCC displays region-specific biomechanical properties to accommodate its biological functions, exhibiting higher stiffness at the anterior region to withstand force-bearing jaw movement, while providing greater cushioning and compliance at the posterior region to absorb mechanical load and support the residence of a potentially stem/progenitor cell niche at the posterior TMJ.

## Materials and Methods

### Animal Strains

Animal experiments were approved by the Institutional Animal Care and Use Committee at the University of Southern California. Both male and female mice were examined as reported in the figure legends.

### Analyses of Mice

Histological analysis, immunostaining, BrdU/EdU incorporation assay, and nanoindentation analysis (UNHT^3^ Bio Nanoindenter, Anton Parr Inc.) were applied to the TMJ and knee joint of postnatal mice. Please refer to the Supplemental Appendix for details.

### Statistics

Statistical analyses were performed in GraphPad Prism 11.0.0 (GraphPad Software Inc, San Diego, CA). Two-tailed Student’s t-test or Welch’s t-test and One-way ANOVA followed by Tukey’s multiple comparison were applied as appropriate, with a significance level at *P* < 0.05.

## Results

### MCC exhibits drastic structural differences from AC at occlusion establishment

Histological analysis of the TMJ and the knee joint was performed on P1, P21, 3-month, and 10-month mice using sagittal sections. At P1, both MCC and AC showed strong proteoglycan staining, with a unique perichondrium layer covering the MCC (Fig. 1A-B’, yellow arrows). At P21, when occlusion is established in mice, we observed drastic structural differences between MCC and AC. MCC at the center of the TMJ condyle was nearly twice as thick as the tibial AC and displayed four distinct zonal layers: superficial, polymorphic, flattened chondrocytes, and hypertrophic chondrocytes (Fig. 1A’, a, b, c, d) (Xu et al. 2022). MCC showed limited proteoglycan staining in its fibrous layer, which comprises the superficial and polymorphic zones (Fig. 1A’, a and b), and strong staining in the cartilaginous layer, formed by flattened chondrocytes and hypertrophic chondrocytes (Fig. 1A’, c and d). In contrast, AC was organized into three proteoglycan-rich zones: superficial, intermediate, and deep zones (Fig. 1B’, e, f, g). By 3 months, MCC exhibited reduced proteoglycan intensity and less distinct zonal boundaries (Fig. 1A’, B’), yet remained thicker than AC (Fig. 1C). By 10 months, MCC displayed a pre-OA-like cartilage degeneration phenotype, including more severe proteoglycan loss, reduced cellularity, decreased cartilage thickness, and mild surface erosions (Fig. 1A’, black arrow), while AC remained largely normal (Fig. 1B’). These histological and structural changes were consistent across sexes (Fig. 1C).

**Figure 1.**
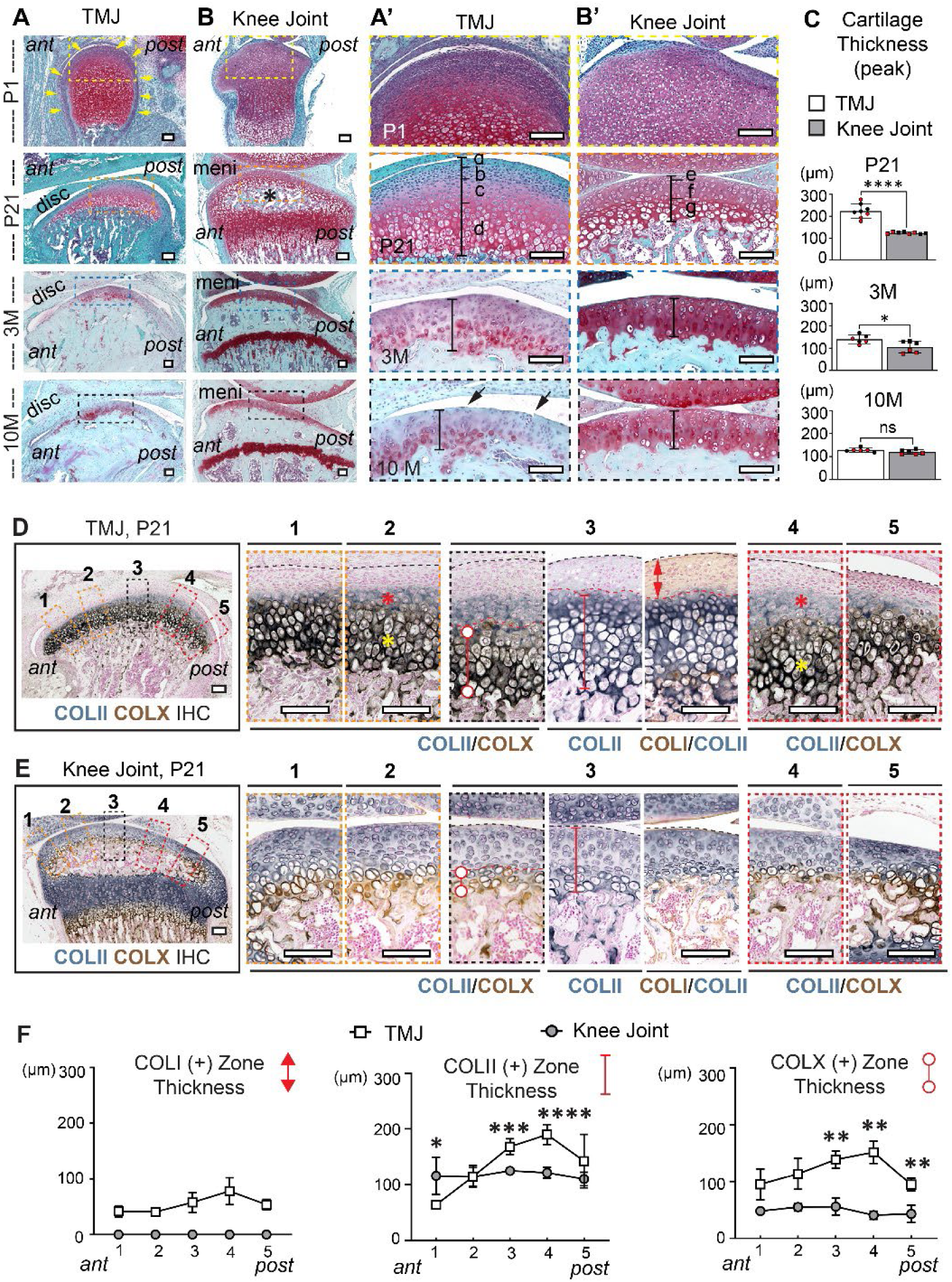
The structure and extracellular matrix composition differ between MCC in the TMJ and AC in the knee joint. **(A, B)** Safranin O/Fast Green staining of TMJ and knee joint sagittal sections at P1, P21, 3 months (3M), and 10 months (10M). MCC and AC are shown in higher magnification in (A’) and (B’). The perichondrium covering MCC is indicated with yellow arrows. The secondary ossification center of the tibia is indicated with a black asterisk. Zonal layers in MCC are indicated with a, b, c, and d, and in AC are indicated with e, f, and g in panel (A’-B’). Quantification of the cartilage thickness at the peak of the TMJ condyle and center of the tibia (black segment) is shown in **(C)**. n=8 for P1, n=6 for P21, 3M, and 12M. An equal number of male and female animals were analyzed. Bars are plotted by means ± SD. Female animals are indicated with red data points. Two-tailed t-test. \**P*<0.05, \*\*\*\**P*<0.0001, ns: not significant. **(D, E)** Immunohistochemistry (IHC) analyses of COLII, COLX, and COLI on TMJ (D) and knee joint (E) at P21. The regions from anterior to posterior of the joints are labeled with 1, 2, 3, 4, and 5. The flattened chondrocytes in MCC are indicated with red asterisks, and the hypertrophic chondrocytes in MCC are indicated with yellow asterisks. The thickness of COLI (+), COLII (+), and COLX (+) zones (red segments) from the anterior to posterior regions of the TMJ and knee joint is quantified in **(F)**. Symbols are plotted with means ± SD. n=6 (3 males, 3 females). ant: anterior; post: posterior; disc: articular disc; meni: meniscus; a: superficial layer; b: polymorphic layer; c: flatten chondrocyte layer; d: hypertrophic chondrocyte layer; e: superficial zone; f: intermediate zone; g: hypertrophic zone. Scalebar: 100μm.

### MCC exhibits a distinct extracellular matrix composition

To assess the regional expression pattern of the extracellular matrix (ECM) components, we analyzed five anterior-to-posterior areas (200μm each) in P21 TMJ and knee joints (Fig. 1D, E). Immunohistochemistry (IHC) analyses of type II collagen (COLII), type X collagen (COLX), type I collagen (COLI), and type III collagen (COLIII) revealed that MCC contains a fibrous layer that is COLI (+)/COLIII (+) but COLII (-) (Fig. 1D and Appendix Fig. 1A, E), and a cartilaginous layer composed of COLII (+)/COLX (-) flattened chondrocytes (Fig. 1D, red asterisk) and COLII (+)/COLX (+) hypertrophic chondrocytes (Fig. 1D, yellow asterisk). Both the fibrous and cartilaginous layers were thickest in the peak-to-posterior MCC (Fig. 1F). In contrast, AC is entirely COLII (+) hyaline cartilage, with a thin layer of COLX (+) hypertrophic chondrocytes in the deep zone (Fig. 1E and Appendix Fig. 1B), and the zonal thickness is uniform across the joint (Fig. 1F). Notably, MCC displayed a markedly thicker COLX (+) zone than AC, especially at the peak-to-posterior regions (Fig. 1F). We also noticed that lubricin (PRG4) expression is enriched in the superficial to intermediate zones of AC, and is more broadly expressed in the superficial, polymorphic, and flattened chondrocyte zones of MCC at P21 (Appendix Fig. 1C, D).

### MCC exhibits a region-specific enrichment of key transcriptional factors

Sex determining region Y-box 9 (SOX9) and runt-related transcription factor 2 (RUNX2), key transcription factors that regulate chondrogenesis and osteogenesis, have been implicated in TMJ development and homeostasis (Hinton et al. 2015; Shibata et al. 2006; Tsutsumi-Arai et al. 2024; Xu et al. 2023). We co-stained COLII with RUNX2 or SOX9 in P21 mice and demonstrated region-specific enrichment of SOX9 and RUNX2 in MCC. Strong SOX9 expression was observed in both the COLII (-) fibrous layer and the COLII (+) cartilaginous layer of MCC (Fig. 2A-A”). A higher percentage of SOX9 (+) cells was present in the posterior region (58.3 ± 22.1%) compared to the anterior region (3.9 ± 2.1%) within the fibrous layer, while its distribution was relatively consistent in the cartilaginous layer (60.7% to 72.6%) (Fig. 2A-A’’, and C). RUNX2 also displayed a higher enrichment in the posterior MCC (72.2 ± 15.5%) than in the anterior region (3.4 ± 0.96%) within the fibrous layer, but was distributed at a comparable level across the cartilaginous layer (65.1% to 82.5%) (Fig. 2D-D”, and F). In AC, SOX9 was highly expressed across all regions (Fig. 2B, B’), while RUNX2 was mainly expressed in the hypertrophic chondrocytes and subchondral bone (Fig. 2E, E’).

**Figure 2.**
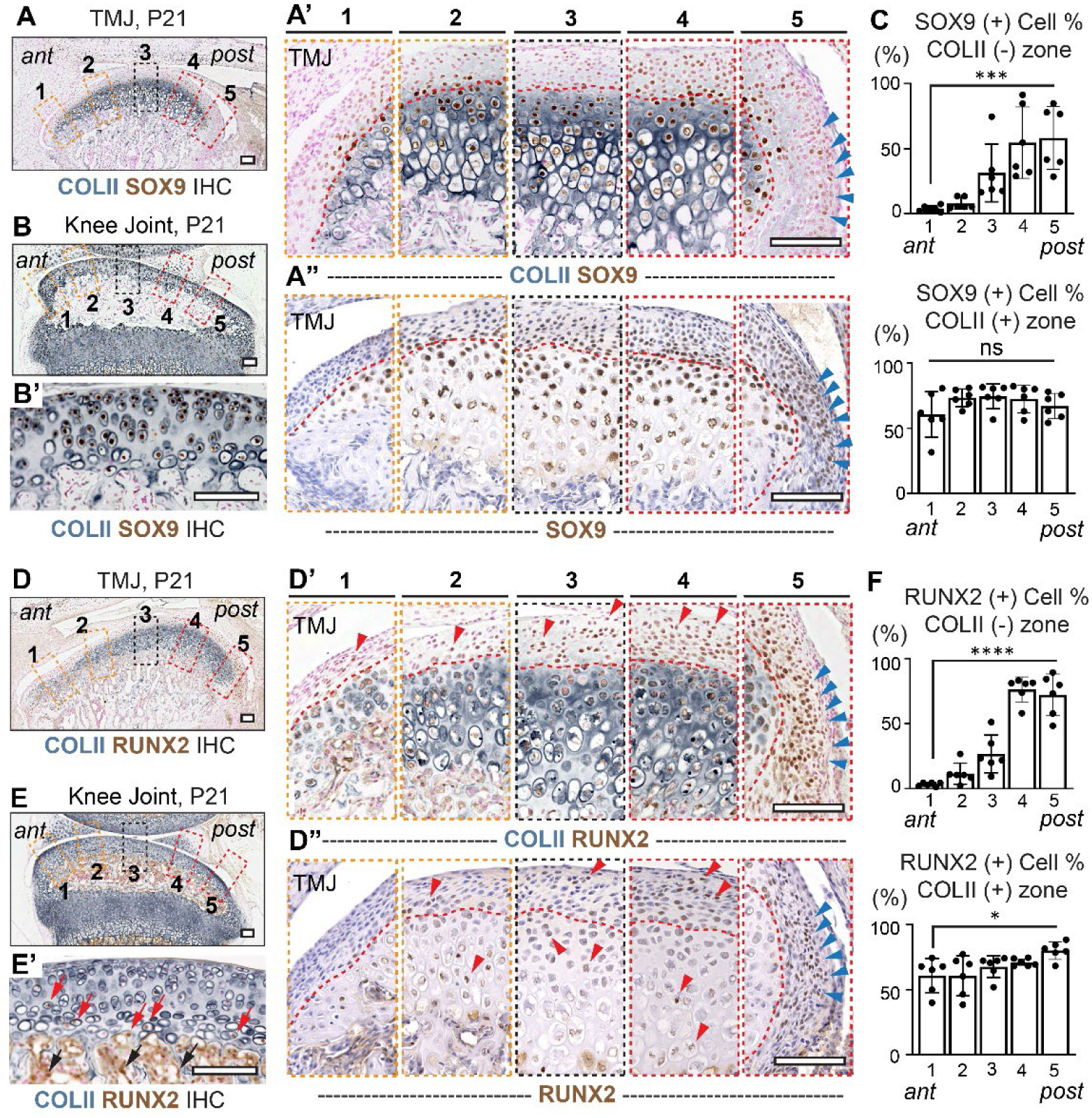
MCC exhibits a region-specific enrichment of SOX9-positive and RUNX2-positive cells at the posterior TMJ. **(A-B’)** IHC analyses of COLII and SOX9 on TMJ (A-A’’) and knee joint (B, B’) at P21. The regions from anterior to posterior of the joints are labeled with 1, 2, 3, 4, and 5. Higher magnification images of MCC are shown in A’ and A’’. Red dot lines indicate the boundaries between the COLII (-) and COLII (+) zones in MCC. SOX9 (+) cells at the posterior region of TMJ are labeled with blue arrowheads. The percentage of SOX9 (+) cells in the COLII (-) and the COLII (+) zones of the MCC is quantified in **(C)**. n=6 (3 males, 3 females). Bars are plotted by means ± SD. One-way ANOVA followed by Tukey’s multiple comparison test. \*\*\**P*<0.001, ns: not significant. **(D-E’)** IHC analyses of COLII and RUNX2 on TMJ (D-D’’) and knee joint (E, E’) at P21. Higher magnification images of MCC are shown in D’ and D’’. Red dot lines indicate the boundaries between the COLII (-) and COLII (+) zones in the MCC of TMJ. RUNX2 (+) cells in the anterior and peak of the MCC are indicated with red arrowheads, and RUNX2 (+) cells in the posterior region of TMJ are indicated with blue arrowheads. RUNX (+) cells in the hypertrophic chondrocytes of the AC and subchondral bone are labeled with red and black arrows, respectively, in (D’ and D’’). The percentage of RUNX2 (+) cells in the COLII (-) and COLII (+) zones of the MCC is quantified in **(F)**. n=6 (3 males, 3 females). Bars are plotted by means ± SD. One-way ANOVA followed by Tukey’s multiple comparison test. \**P*<0.05, \*\*\*\**P*<0.0001, ns: not significant. ant: anterior; post: posterior. Scalebar: 100μm.

We next examined the expression pattern of scleraxis (SCX), the crucial transcription factor for tendon and ligament development that also regulates TMJ formation (Blitz et al. 2009; Bobzin et al. 2021; Roberts et al. 2019). SCX expression was robust in the fibrous layer and the flattened chondrocytes of MCC at P1, P21, and 2 months, especially in the posterior MCC (Appendix Fig. 2A-C), with low-level expression in AC (Appendix Fig. 2D). Lineage tracing with the *ScxCre; Rosa-mTmG^f/+^* reporter mice revealed GFP (+) cell patches extending from the MCC surface into the polymorphic and the flattened chondrocyte zones at P21 (Appendix Fig. 2E). By 2 months, the entire MCC is GFP (+) (Appendix Fig. 2F), indicating a significant contribution of the *Scx*-expressing cells to postnatal formation of MCC, consistent with previous studies (Ma et al. 2021; Roberts et al. 2019).

### The progenitor-like, slow-cycling cells are enriched in the posterior TMJ

To investigate the cell proliferation profiles in postnatal TMJ, we performed BrdU/EdU double incorporation analyses. BrdU was administered from P6-P10 and chased until P21 to label the progenitor-like, slow-cycling cells, and EdU was administered 4 hours before tissue harvesting to label actively proliferative cells (Fig. 3A). We found that the BrdU (+) label-retaining cells (slow-cycling cells) were mainly located at the MCC surface (Fig. 3B-B’’, yellow arrows), consistent with previous findings (Lei et al. 2022; Tuwatnawanit et al. 2025a). Interestingly, these slow-cycling cells are enriched at the posterior TMJ (Fig. 3B’’’, yellow arrows), which overlaps with the SOX9, RUNX2, and SCX-enriched area (Fig. 2A-A’’, D-D”, and Appendix Fig. 2A-C). The EdU (+) actively proliferative cells, on the other hand, were mainly observed in the polymorphic cells of the MCC (Fig. 3B-B’’, white arrows). In the AC, BrdU (+) cells were mainly located at the superficial zone, while the EdU (+) cells were almost undetectable (Fig. 3C’), suggesting that the AC cells have reached homeostasis at P21. The growth plate is a specialized structure responsible for the longitudinal growth of the long bone, which houses slow-cycling cells in the resting zone (Fig. 3C’’, white arrows) and actively proliferative cells in the proliferative zone (Fig. 3C’’, yellow arrows), as previously reported (Bian et al. 2024). Next, we administered EdU from P6-P10 and chased until 4.5 months to trace the progenitor-like, label-retaining cells (Fig. 3D). We found that the EdU (+) label-retaining cells (slow-cycling cells) were only visible at the posterior region of the TMJ (Fig. 3E, yellow arrows) but were diminished from the MCC peak (Fig. 3E). These results confirm that the polymorphic zone of MCC contains highly proliferative cells, while the surface of MCC, especially the posterior region, houses progenitor-like, slow-cycling cells that persist for a longer time during the postnatal maintenance of TMJ.

**Figure 3.**
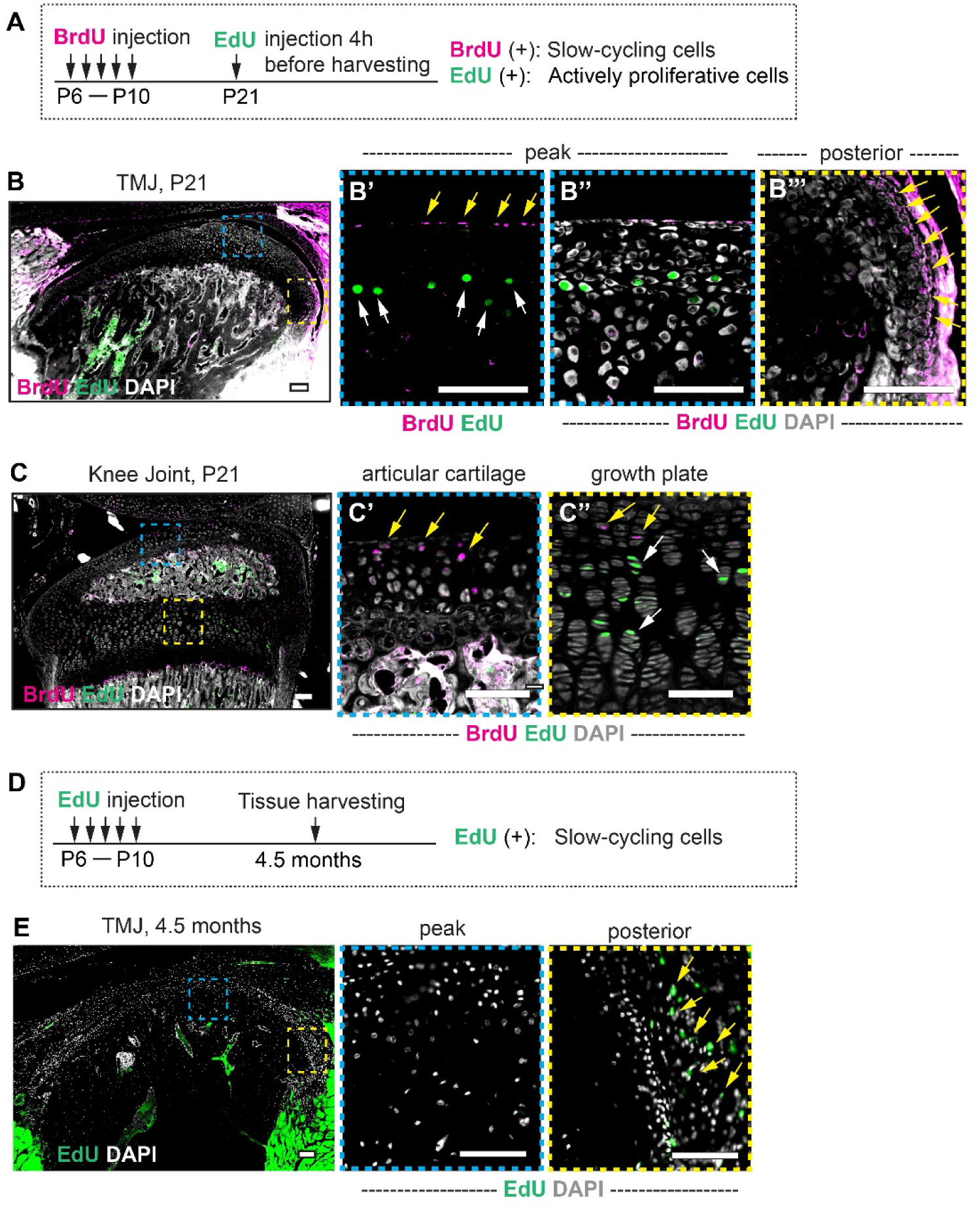
The progenitor-like, slow-cycling cells within the fibrous layer of the MCC are enriched in the posterior region of the TMJ. **(A)** The scheme used for the BrdU and EdU double incorporation assay to label slow-cycling cells and actively proliferative cells. **(B, C)** BrdU and EdU staining performed on P21 TMJ and knee joint sections. The peak and posterior region of the TMJ are shown in higher magnification in (B’-B’’’), and the AC and growth plate in the tibia are shown in (C’, C”). Slow-cycling cells in the peak and posterior TMJ are indicated with yellow arrows in (B’) and (B’’’), and actively proliferative cells in MCC are indicated with green arrows in (B’). Slow-cycling cells in AC and the resting zone of the growth plate are indicated with yellow arrows in (C’, C’’), and actively proliferative cells in the growth plate are indicated with white arrows in (C”). n=4 (2 males, 2 females). **(D)** The scheme used for the EdU incorporation assay to trace the slow-cycling, label-retaining cells. **(E)** EdU staining performed on 4.5-month TMJ sections. EdU (+) slow-cycling cells are indicated with yellow arrows. n=4 (2 males, 2 females). Scalebar: 100μm.

### MCC exhibits a lower and regionally gradient elastic modulus compared to AC

Next, we compared the biomechanical properties between MCC and AC utilizing a high-resolution, automated nanoindentation mapping technique. Intact cartilage of MCC and AC was tested with a consistent force to mimic biomechanical impact under low-level physiological loading (Fig. 4A) (Lin and Horkay 2008), followed by assessment of the force-displacement curve in P21 mice (Fig. 4B). The Young’s modulus/elastic modulus (E_IT_), which measures the resistance of cartilaginous tissue to elastic deformation, was calculated for each measurement position (Kuroda et al. 2009; Petitjean et al. 2023). To facilitate analysis, MCC were separated into three regions: anterior, peak, and posterior; while AC (both the lateral and medial tibial AC) were separated into three zones: center (zone 1, non-load-bearing), anterior peripheral (zone 2, load-bearing), and posterior peripheral (zone 3, load-bearing) (Fig. 4C). We found that the E_IT_ of MCC was significantly lower than AC (Fig. 4E). Within MCC, the anterior region showed the highest E_IT_ (0.281 to 0.807 MPa) and the posterior showed the lowest (0.168 to 0.430 MPa) (Fig. 4D, F). Within AC, load-bearing regions showed higher E_IT_ (zones 2 and 3, 0.1 to 5.539 MPa) than the non-load-bearing region (zone 1, 0.081 to 2.656 MPa) in both lateral and medial AC (Fig. 4D, G), which is consistent with previous findings (Briant et al. 2015). Since we optimized nanoindentation to measure MCC and AC with the same protocol and parameters, the biomechanical features between these two tissues can be directly compared. We found that even the stiffest anterior region of MCC exhibits a 67.44% reduction in E_IT_ compared to the load-bearing regions of AC (Fig. 4F, G). We observed consistent biomechanical features across sexes, so we further analyzed male and female MCC in one cohort (n = 10, 5 males, 5 females). Spatial profiling of MCC’s mechanical properties using a topographical heatmap (Fig. 5A) revealed that the mean E_IT_ in the anterior region (0.523 ± 0.112 MPa) was significantly higher than the peak and posterior regions (0.373 ± 0.095 MPa and 0.293 ± 0.072 MPa, respectively) (Fig. 5B). The elastic work of indentation (W_el_), which reflects the recovered portion of the total mechanical work done during nanoindentation following indenter removal from the tissue (Fischer-Cripps 2011), shows an opposite profile to the E_IT_ (Fig. 5C). These data suggest that the anterior MCC is stiffer and more resistant to mechanical deformities, while the posterior MCC is more compliant with greater capability for force absorption.

**Figure 4.**
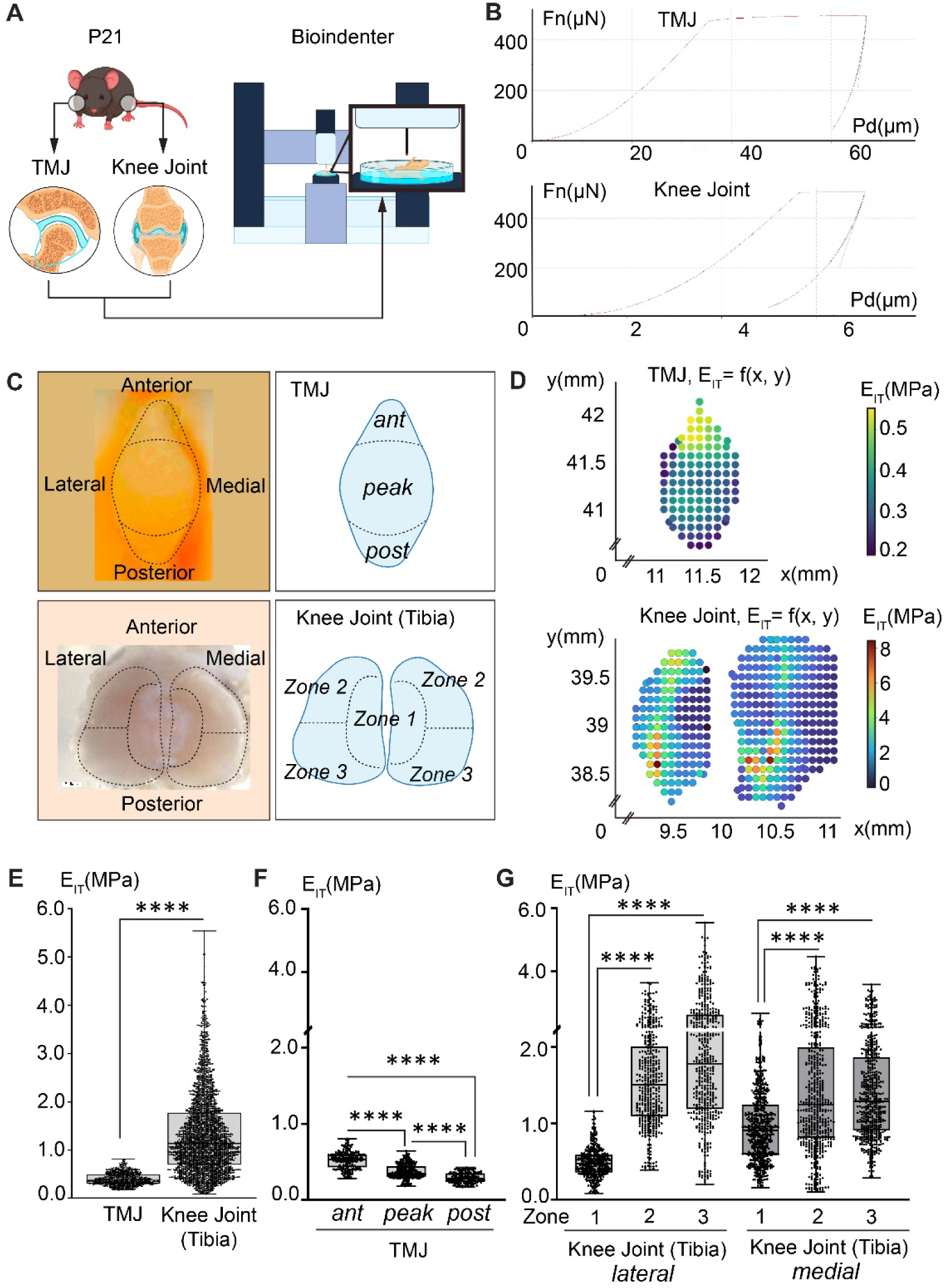
MCC exhibits a lower and regionally gradient elastic modulus compared to AC. **(A)** Schematic of the biomechanical testing using a Bio Nanoindenter. **(C)** Top views and the schematics of the zonal regions of the TMJ condyle and tibia plateau of P21 mice. **(B)** Representative force-displacement curve generated with MCC of the TMJ and AC of the knee joint. **(D)** Representative heatmap of the elastic modulus (E_IT_) generated on the intact cartilage surface of the MCC and AC. Each dot represents one nanoindentation measurement position. **(E)** MCC exhibits a significantly lower E_IT_ compared with the AC. Boxes and whiskers are plotted with all measurements shown from Min to Max. Each dot represents one nanoindentation measurement data point. n=784 for MCC in TMJ and n=3028 for AC in the tibia of the knee joint. Two-tailed Welch’s t-test. \*\*\*\**P*<0.0001. **(F, G)** MCC exhibits the highest E_IT_ in the anterior region and the lowest E_IT_ in the posterior region (F), while AC exhibits higher E_IT_ in the load-bearing region (zones 2 and 3) and lower E_IT_ in the non-load-bearing region (zone 1). Boxes and whiskers are plotted with all data shown from Min to Max. Each dot represents one nanoindentation measurement data point. In TMJ, n=180 for anterior, n=361 for peak, and n=203 for posterior. For the lateral tibia of the knee joint, n= 395 for zone 1, n= 423 for zone 2, and n=476 for zone 3; for the medial tibia of the knee joint, n= 616 for zone 1, n=548 for zone 2, and n=571 for zone 3. One-way ANOVA followed by Tukey’s multiple comparison test. \*\*\*\**P*<0.0001,

**Figure 5.**
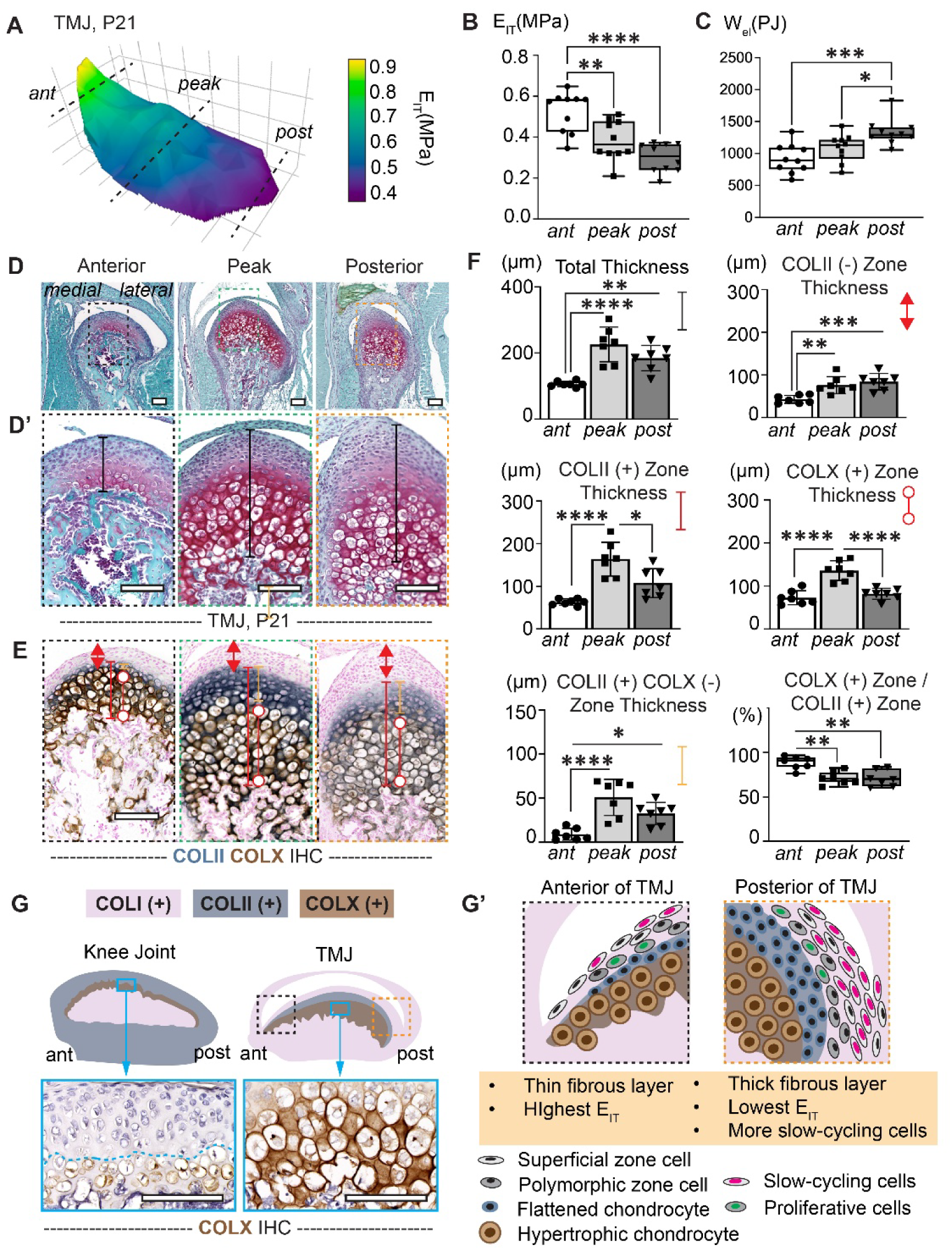
Regional biomechanical properties of MCC correlate with ECM architecture. **(A-C)** 3D topographical heatmap (A) and quantification of E_IT_ (B) and W_el_ (C) across the TMJ condyle. n=10 (5 males, 5 females). **(D-D’)** Safranin O/Fast Green staining of TMJ coronal sections at P21 from the anterior, peak, and posterior planes, as indicated in (A). Higher magnifications of MCC are shown in (D’). **(E)** IHC analyses of COLII and COLX on MCC at P21. The thickness of COLII (-), COLII (+), and COLX (+) zones (red segments) and COLII(+)/COLX(-) zone (orange segment) are indicated. **(F)** The total thickness of the MCC cartilage, the thickness of the COLII (-), COLII (+), COLX (+), and the COLII (+)/COLX (-) zones, and the ratio between the COLX (+) and COLII (+) zones are shown. n=7 (4 males, 3 females). Bars are plotted with means ± SD. One-way ANOVA followed by Tukey’s multiple comparison test. \**P*<0.05, \*\**P*<0.01, \*\*\**P*<0.001, \*\*\*\**P*<0.0001. **(G, G’)** Illustrations of the ECM composition between AC in the knee joint and MCC in the TMJ (G), and the MCC architecture between the anterior and posterior regions of the TMJ (G’). Scalebar: 100μm.

### Regional biomechanical properties of MCC correlate with ECM architecture

To further dissect the association between the spatial distribution of biomechanical properties and MCC structures, we performed histological analysis and IHC staining of COLII and COLX on coronal sections of P21 TMJ at the anterior, peak, and posterior planes (Fig. 5A, D-E). Quantification of total cartilage thickness reveals that the anterior MCC has thinner cartilage than the peak and posterior regions (Fig. 5D’ and F). IHC analysis shows the thinnest COLII (-) fibrous zone and the thinnest COLII (+)/COLX (-) flattened chondrocytes zone at the anterior MCC (Fig. 5E, F), which is negatively associated with the E_IT_ (Fig. 5B). Though the highest thickness of COLII (+) and COLX (+) zones were observed at the peak region (Fig. 5E, F), the anterior region exhibits the highest ratio of the COLX (+) zone thickness to the COLII (+) zone thickness (Fig. 5E), which is positively correlated with the E_IT_ (Fig. 5B). These data suggest that MCC stiffness may be closely correlated with the cartilage architecture and ECM composition. A higher E_IT_ may be associated with a thinner fibrous layer and a higher percentage of COLX (+) ECM in the cartilaginous layer of MCC.

## Discussion

### MCC shows distinct maturation and degeneration patterns compared to AC

Our study demonstrates different maturation and degeneration patterns for MCC and AC in mice. MCC is markedly thicker than AC at the establishment of occlusion, which is a key milestone in TMJ maturation associated with solid food intake, but undergoes earlier and more pronounced degeneration in pre-aged mice (10 months), whereas AC typically develops spontaneous OA-like changes much later, between 17 and 24 months (Geraghty et al. 2023). These findings align with prior reports and mirror human epidemiology, where TMJ OA manifests earlier in life than knee OA (Bielajew et al. 2021; Chen et al. 2020; Tang et al. 2025). The earlier degeneration of MCC may arise from its dynamic remodeling nature, which relies on the replenishment and renewal of the fibrocartilage stem cells (FCSCs) within the fibrous layer (Tuwatnawanit et al. 2025a). Age-related loss of the fibrous layer and exhaustion of the FCSC likely compromise regenerative capacity, leaving the underlying cartilaginous layer more vulnerable to biomechanical stress (Kuroda et al. 2009). These insights are consistent with the clinical observations that TMJ OA has a multifactorial and complex pathoetiology, where mechanical overloading alone cannot account for the full spectrum of disease phenotypes (Clark et al. 2025).

### The biomechanical properties of MCC may support a stem/progenitor cell niche

Distinct populations of fibrocartilage stem cells (FCSCs) have been identified within MCCs from mice, rats, rabbits, and humans (Bi et al. 2020; Embree et al. 2016; Tuwatnawanit et al. 2025a; Tuwatnawanit et al. 2025b). These progenitor/stem cells have been localized to the superficial zone of MCC using the BrdU/EdU retention assay, as label-retaining cells (LRCs) represent a hallmark of stem cells (Embree et al. 2016; Tuwatnawanit et al. 2025b; Wang et al. 2024). Our studies also identified the residence of these LRCs/slow-cycling cells at the MCC surface and further demonstrated their enrichment at the posterior region of TMJ. This region overlaps with elevated expression of key transcriptional factors for mesenchymal lineage differentiation (SOX9, RUNX2, and SCX) and is maintained into later adulthood. These observations collectively suggest that the posterior region of TMJ may harbor a long-lasting stem/progenitor cell niche.

The location of this niche is closely associated with the biomechanical properties of MCC. Our data revealed a distinct regional distribution of the biomechanical properties of MCC, stiffest in the anterior region, followed by the peak and posterior regions, consistent with previous large animal studies utilizing atomic force microscopy nanoindentation in the rabbit condyle (Hu et al. 2001) and dynamic indentation with pig condyles (Tanaka et al. 2006). These findings verify the mouse model as a valuable system to study biomechanical properties of the TMJ.

During mastication in humans and mice, the anterior MCC is exposed to compressive and shear forces, with the anterior ridge of the mandible bearing the largest stress and the highest load-induced strain (Miyawaki et al. 2001; Tsouknidas et al. 2017). The biochemical stress profile is known to regulate anterior MCC remodeling, as mice fed with hard food displayed a thinner fibrous layer in the anterior region than mice fed with a soft diet, but the posterior region was insensitive to diet-induced remodeling (Uptegrove et al. 2025). The anterior MCC in humans is also a major load-bearing zone during clenching (Nishio et al. 2009). Meanwhile, the posterior TMJ is exposed to tensile forces during jaw movement (Murphy et al. 2013; Singh and Detamore 2008), which may be functionally linked to the high content of progenitor-like cells, as cyclic tensile strain modulates progenitor cell differentiation towards the osteogenic and chondrogenic fates (Gilbert et al. 2021; Zhang et al. 2024). Taken together, the anterior MCC bears compressive and shear forces to facilitate the gliding motion of the TMJ (Ruggiero et al. 2015), while the posterior MCC, with thicker fibrous and hyaline cartilage layers, achieves a force-absorbent function that dissipates shocks during speech and biting to protect the brain and other cranial structures from damage (Porter et al. 2025), and may support the residence of a potential stem/progenitor cell niche (Fig. 5G, G’).

### ECM composition underlies the regional biomechanical properties of MCC

Our study suggests that a higher elastic modulus of MCC is associated with a thinner fibrous layer and a cartilaginous layer with a higher ratio of COLX (+) ECM thickness. Therefore, the biomechanical properties of MCC may reflect a complex interplay of the fibrous layer and the underlying flattened chondrocytes and hypertrophic chondrocytes. Compared with the hypertrophic chondrocytes in AC, which only exist in the deep zone cartilage, the hypertrophic chondrocytes of the MCC are major constituents of the MCC, making up around 90% of the cartilaginous zone thickness at the anterior region and around 70% in the posterior region (Fig. 5E). Moreover, these MCC hypertrophic chondrocytes tend to be more compact with higher cell density, less ECM, and a more intense pericellular matrix in mice (Fig. 5G), similar to the observations in pigs (Porter et al. 2025), suggesting an additional role of the hypertrophic chondrocytes in supporting the viscoelastic properties of the MCC.

In conclusion, this study establishes a comprehensive foundation for comparing the biological and biomechanical properties of MCC and AC in postnatal mice. By integrating unbiased nanoindentation mapping with histological and biochemical analyses, we identify a region-specific structure-function relationship unique to MCC, underscoring the necessity of developing TMJ-specific animal models. This study advances our understanding of the fundamental differences between the TMJ and knee joint, provides important context for interpreting TMJ disease progression, and supports the development of mouse models to study TMJ-specific pathologies.

## Author Contributions

JH: Contributed to data acquisition, data analysis, data interpretation, and drafted and critically revised the manuscript. HCF and JG: Contributed to data acquisition, data analysis, data interpretation, and critically revised the manuscript. JRL, MV, LV, and JBE: Contributed to data acquisition, data analysis, and critically revised the manuscript. FB, HKJ, and CL: Contributed to data acquisition and critically revised the manuscript. PK: Contributed to data analysis, data interpretation, and critically revised the manuscript. GTC, AVM, AEM, and JC: Contributed to data interpretation and critically revised the manuscript. JX and ZL: Contributed to conceptualization, design, data acquisition, data interpretation, supervision, project administration, and drafted and critically revised the manuscript. All authors gave final approval and agreed to be accountable for all aspects of the work.

## Acknowledgments

We thank Dr. Ronen Schweitzer for sharing the *ScxCre* mouse strain. We thank Mr. Thach-Vu Ho at the Center for Craniofacial Molecular Biology at USC for technical support.

## Supplemental Appendix

## Appendix Materials and Methods

### Animal Strains

All the animal experiments were approved by the Institutional Animal Care and Use Committee at the University of Southern California (Protocols 21421 and 21688). Mice used in this study include C57B/6J (JAX: 000664, The Jackson Laboratory), *ScxCre* (JAX: 032131, The Jackson Laboratory) (Blitz et al. 2009), and *ROSA-mTmG* reporter (JAX: 007676, The Jackson Laboratory) (Muzumdar et al. 2007). Both male and female mice were examined, with the number and sex reported in the figure legends.

### Analyses of Mice

Histological analyses were performed on P1, P21, 3-month, and 10-month mice. Mouse jaw and hindlimbs that were fixed in 10% neutral-buffered formalin (Sigma HT501128) for 1–3 days, followed by 1–5 days of decalcification in Formic Acid Bone Decalcifier (Immunocal, StatLab UN3412). Tissues were embedded in paraffin and sectioned at 5μm thickness. Safranin O/Fast Green staining was performed following standard protocols (Center for Musculoskeletal Research, University of Rochester).

Immunohistochemistry (IHC) analyses were performed with antigen retrieval (COLII and COLX: 4mg/ml pepsin in 0.01 N HCl solution, 37°C water bath for 10 min; COLI and COLIII: 10μg/ml Proteinase K for 10 minutes at room temperature; SOX9, RUNX2, SCX, and GFP: 10mM Tris-EDTA with 0.05% Triton-X-100, pH 9.0, 75°C water bath for 5 min; PRG4: 100μg/ml hyaluronidase, 37°C water bath for 10 min), and incubation with the following primary antibodies: anti-COLII (Thermo Scientific MS235, 1:200), anti-COLX (Abclonal, A18604, 1:100), anti-COLI (abcam, ab138492, 1:1000), anti-COLIII (abcam, ab7778, 1:1000), anti-SOX9 (Millipore/Sigma-Aldrich, AB5535, 1:400), anti-RUNX2 (Abclonal, A2851, 1:200), anti-SCX (Invitrogen, PA5-115874, 1:200), anti-GFP (ABclonal, AE011, 1:400), and anti-Lubricin (PRG4) (abcam, ab28484, 1:400). The corresponding mouse IgG or rabbit IgG controls were also performed. Targets were detected via secondary antibody incubation (Vector Laboratories PK-6200 for IHC, Invitrogen A-11034 for IF) and colorimetric developmental methodologies (DAB substrate, Vector Laboratories SK-4105; SG Substrate, Vector Laboratories SK-4700). Nuclei were counterstained with hematoxylin or nuclear fast red.

BrdU/EdU incorporation assays were performed as indicated in the figures. For BrdU/EdU double labeling, BrdU was injected i.p. (100 µg/g) five times from P6 to P10, and EdU was injected i.p. (25µg/g) 4 hours before euthanasia at P21. For EdU chase analysis, EdU was injected i.p. (25 µg/g) five times from P6 to P10, and the animals were harvested at 4.5 months. BrdU was detected using an anti-BrdU antibody (Thermo Fisher B35128, 1:200), followed by a secondary antibody Alexa Fluor 594 (Invitrogen A32742, 1:1500). EdU was detected using the Click-iT EdU Cell Proliferation Kit (Invitrogen C10337) according to the manufacturer’s instructions.

Stained sections were photographed using a Keyence BZX810 microscope. Cartilage thickness and cell number are quantified on the high-resolution bright field or fluorescence images with ImageJ (https://imagej.net/). Five areas with a width of 200 μm were selected along the anterior to posterior of the TMJ and the knee joint for detailed analysis. Area 3 was located at the peak of the TMJ or the center of the AC, while areas 1/2 and 4/5 were located at the anterior or posterior of the joint, respectively.

### Automated Nanoindentation

Mandibles and hindlimbs of P21 C57B/6J mice were dissected and stored at −20 °C in PBS-saturated gauze until ready for biomechanical testing. On the testing day, samples were thawed at room temperature and mounted for testing. Specifically, the body of the mandible was immersed in a layer of dental wax in a 35mm dish and held by the neck area using forceps until the wax hardened (Appendix Fig. 3A). The tibia, with the plateaus facing up, was inserted into a 10ul pipette tip (VWR, USA) pre-glued to the center of a 35mm diameter culture dish and further fixed with polyurethane adhesive (Gorilla gel glue) to prevent any displacement during indentation (Appendix Fig. 3B). The samples were then immersed in PBS and inspected under a dissection microscope to ensure no glue was on or around the cartilage surface. The samples were fully immersed in PBS during the whole test to ensure hydration of the tissue during measurement. The 35mm dish is securely mounted on the testing stage of UNHT^3^ Bio Nanoindenter (Anton Parr Inc.) equipped with an active vibration isolation unit to mitigate the influence of external vibration that affects nanoindentation analysis. The measurement was conducted using a spherical indenter with a 500 μm radius, which allows local characterization while at the same time reducing the effect of cartilage heterogeneity and surface unevenness. For the automated indentation of tibial AC and MCC (N = 5 female and 5 male mice), a standardized rectangular mapping matrix with 100nm spacing was superimposed on the image of the cartilage surface, which included 100 to 150 measurement positions for the lateral and 150 to 200 for the medial tibial condyle, and 100 to 150 measurement positions for the TMJ mandibular condyle. Automated indentation at each predefined position started with a sample surface identification step through the adjust depth offset (ADO) procedure, followed by indentation with a maximum load of 200μN and a hold period of 60 seconds at maximum load (Fmax) to observe the time-dependent behavior of the cartilage under constant load. The loading and unloading times were set to 30 seconds. The force-time and displacement-time were recorded for the identification of the approach, loading phase, pause, and unloading phase, and the force-displacement curve was analyzed. In all experiments, Young’s modulus or elastic modulus (E_IT_) and elastic work of indentation (W_el_) were calculated by the Oliver and Pharr fitting to the unloading portion of the force-displacement curve. Each parameter (i.e., E_IT_, W_el_) was compared between lateral AC, medial AC, and MCC, in sub-regions as indicated in Figure 4. Each region contains at least 100 positions of measurement. Outliers were identified using the ROUT method (Robust regression and Outlier removal) with a coefficient Q set to 1% (GraphPad Prism 9.5.0). Topographical mapping of the MCC biomechanical properties was performed based on E_IT_ and the spatial location of each measurement point.

## Appendix Figures and Figure Legends

**Appendix Figure 1.**
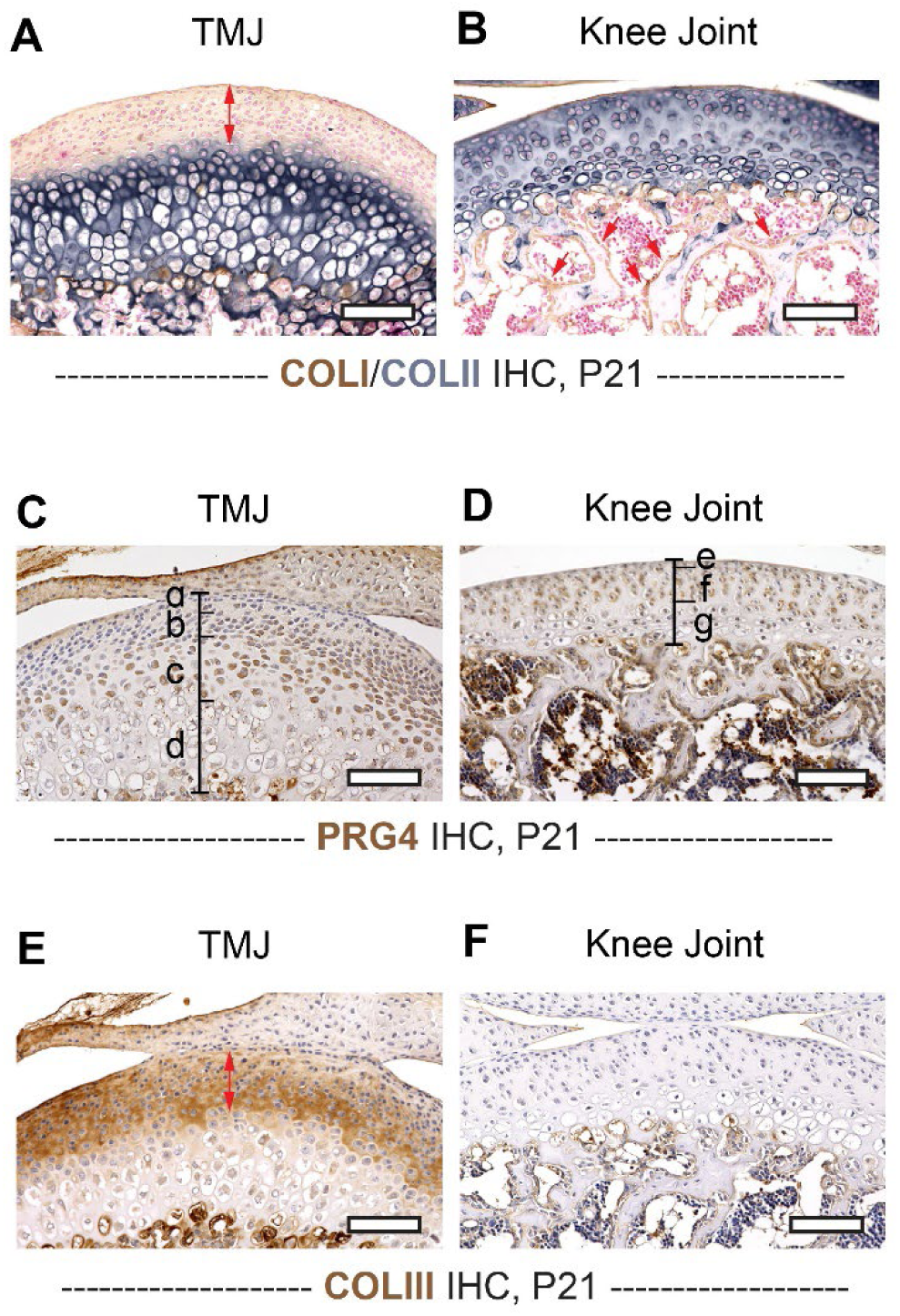
IHC analyses on additional ECM components in MCC and AC. **(A, B)** IHC analyses of COLII and COLI on MCC (A) and AC (B). The COLII (-)/COLI (+) fibrous layer in MCC is indicated with a red segment with double arrowheads. The COLI (+) bone matrix in the subchondral bone of the knee joint is indicated with red arrows. **(C, D)** IHC analyses of PRG4 on MCC (C) and AC (D). The PRG4 (+) cells in MCC are mainly located in the superficial (a), polymorphic (b), and the flattened chondrocytes (c), but not the hypertrophic chondrocytes (d). The PRG4 (+) cells in AC are mainly located in the superficial (e) and the intermediate (f) zones, but not the hypertrophic zone (g). **(E, F)** IHC analyses of COLIII on MCC (E) and AC (F). The COLIII (+) fibrous layer in MCC is indicated with a red segment with double arrowheads. n=6. Scalebar: 100μm.

**Appendix Figure 2.**
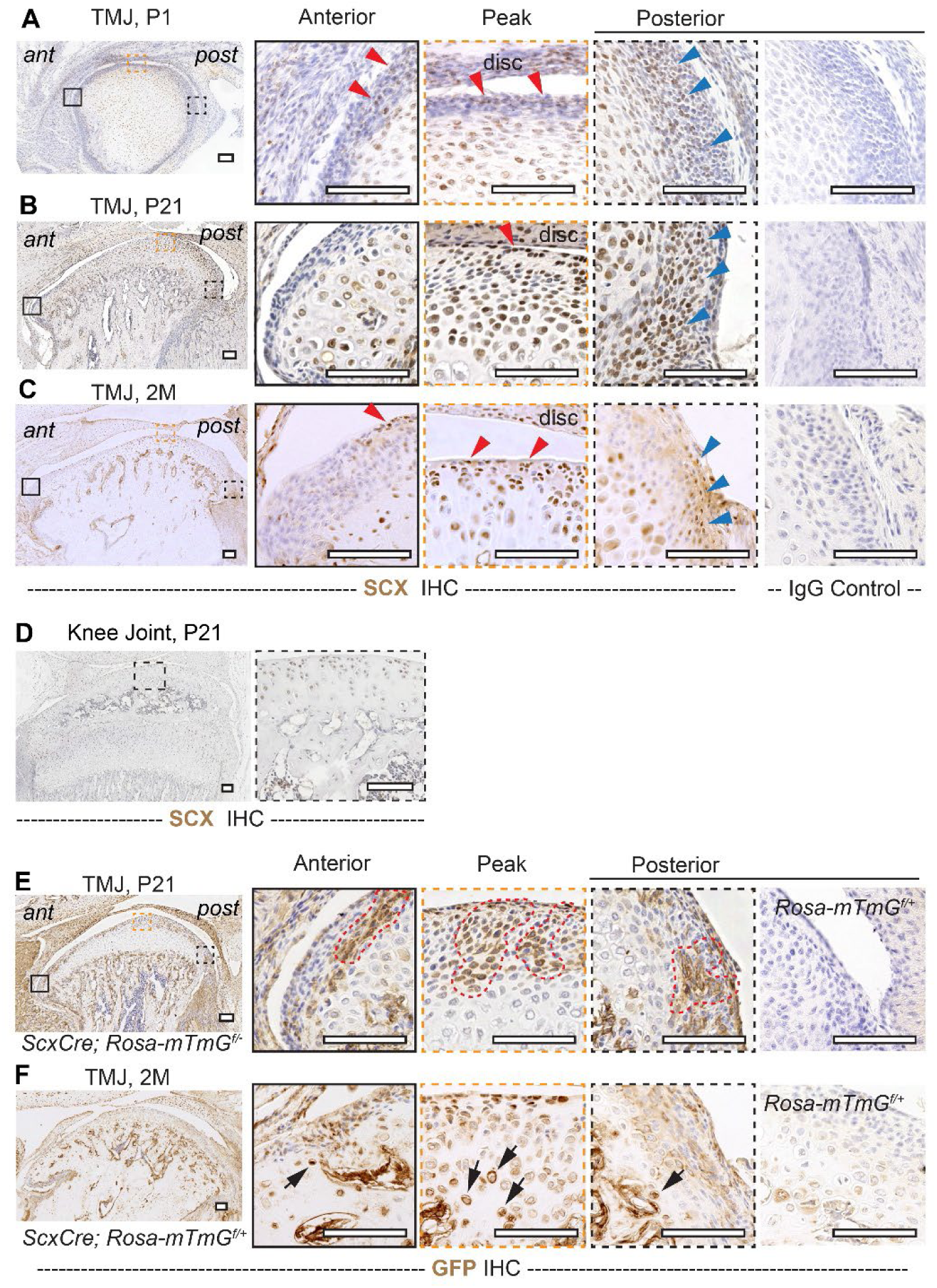
SCX-positive cells contribute to the postnatal development of MCC. **(A-D)** IHC analyses of SCX on wild-type TMJ sagittal sections at P1 (A), P21 (B), and 2 months (2M) (C), and on wild-type knee joint sections at P21 (D). Red arrowheads indicate SCX(+) cells in the fibrous layer of MCC. Highly enriched SCX expression at the posterior regions of the TMJ is indicated with blue arrowheads. Negative IgG control is also shown. n=4. **(E-F)** IHC analysis of GFP on TMJ sections of *ScxCre; Rosa-mTmG^f/+^* mice or *Rosa-mTmG^f/+^* mice at P21 and 2 months (2M). Red dotted lines indicate GFP(+) cell patches in MCC at P21 (E), and black arrows indicate GFP(+) cells in the hypertrophic chondrocytes of the MCC at 2 months (F). n=3. ant: anterior; post: posterior. Scalebar: 100μm.

**Appendix Figure 3.**
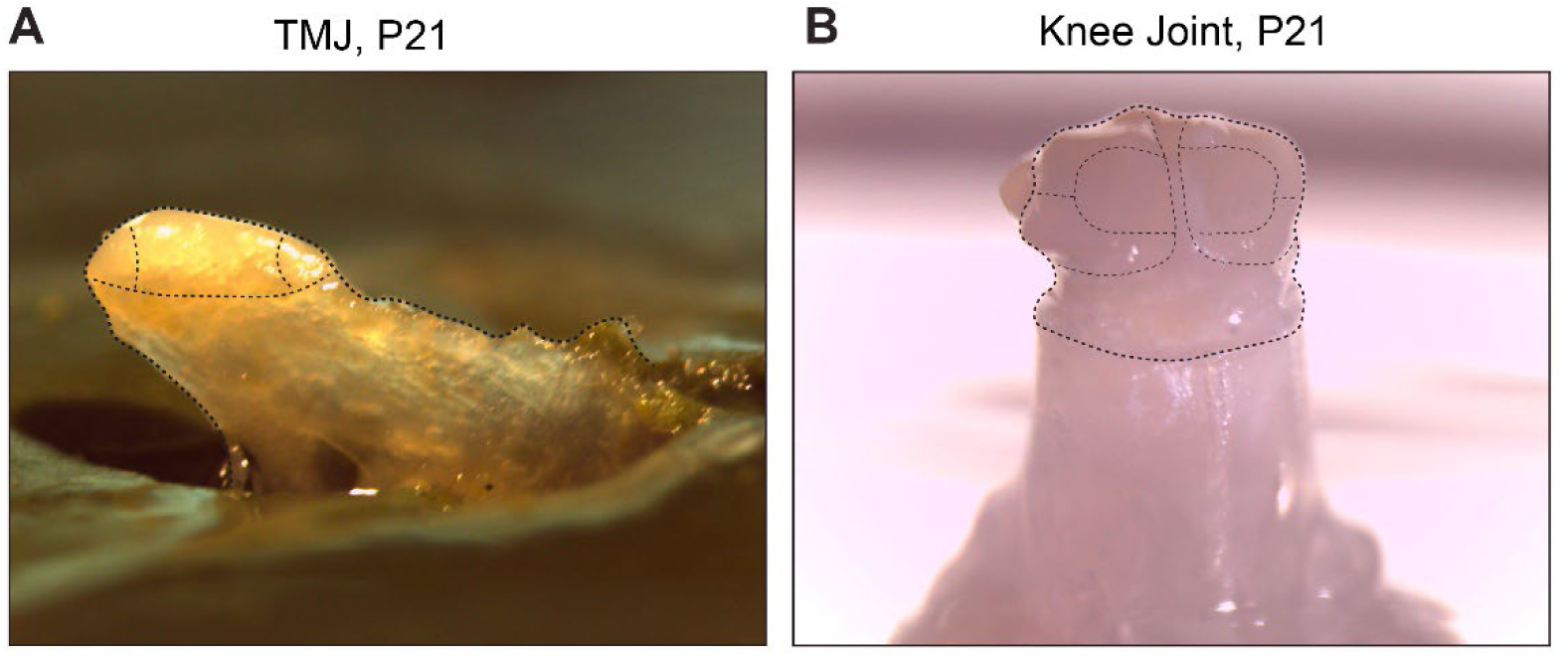
Illustration of the TMJ and knee joints mounted for biomechanical testing. **(A, B)** Representative images of the TMJ (A) and tibia (B) mounted to the center of a culture dish. The analysis regions and zones are indicated with dotted lines.

## Reference

Bi R, Yin Q, Mei J, Chen K, Luo X, Fan Y, Zhu S. 2020. Identification of human temporomandibular joint fibrocartilage stem cells with distinct chondrogenic capacity. Osteoarthritis Cartilage. 28(6):842–852.

Bian F, Hansen V, Feng HC, He J, Chen Y, Feng K, Ebrahimi B, Gray RS, Chai Y, Wu CL et al. 2024. The g protein-coupled receptor adgrg6 maintains mouse growth plate homeostasis through ihh signaling. J Bone Miner Res.

Bielajew BJ, Donahue RP, Espinosa MG, Arzi B, Wang D, Hatcher DC, Paschos NK, Wong MEK, Hu JC, Athanasiou KA. 2021. Knee orthopedics as a template for the temporomandibular joint. Cell Rep Med. 2(5):100241.

Blitz E, Viukov S, Sharir A, Shwartz Y, Galloway JL, Pryce BA, Johnson RL, Tabin CJ, Schweitzer R, Zelzer E. 2009. Bone ridge patterning during musculoskeletal assembly is mediated through scx regulation of bmp4 at the tendon-skeleton junction. Dev Cell. 17(6):861–873.

Bobzin L, Roberts RR, Chen HJ, Crump JG, Merrill AE. 2021. Development and maintenance of tendons and ligaments. Development. 148(8).

Briant P, Bevill S, Andriacchi T. 2015. Cartilage strain distributions are different under the same load in the central and peripheral tibial plateau regions. J Biomech Eng. 137(12):121009.

Chen PJ, Dutra EH, Mehta S, O’Brien MH, Yadav S. 2020. Age-related changes in the cartilage of the temporomandibular joint. Geroscience. 42(3):995–1004.

Clark GT, Vistoso Monreal A, Veas N, Loeb GE. 2025. Artificial intelligence algorithm for real-time diagnostic assist in orofacial pain. J Am Dent Assoc. 156(8):664–673.

Embree MC, Chen M, Pylawka S, Kong D, Iwaoka GM, Kalajzic I, Yao H, Shi C, Sun D, Sheu TJ et al. 2016. Exploiting endogenous fibrocartilage stem cells to regenerate cartilage and repair joint injury. Nat Commun. 7:13073.

Fischer-Cripps AC. 2011. Nanoindentation. Springer New York, NY.

Geraghty T, Obeidat AM, Ishihara S, Wood MJ, Li J, Lopes EBP, Scanzello CR, Griffin TM, Malfait AM, Miller RE. 2023. Age-associated changes in knee osteoarthritis, pain-related behaviors, and dorsal root ganglia immunophenotyping of male and female mice. Arthritis Rheumatol. 75(10):1770–1780.

Gilbert SJ, Bonnet CS, Blain EJ. 2021. Mechanical cues: Bidirectional reciprocity in the extracellular matrix drives mechano-signalling in articular cartilage. Int J Mol Sci. 22(24).

Hinton RJ, Jing J, Feng JQ. 2015. Genetic influences on temporomandibular joint development and growth. Curr Top Dev Biol. 115:85–109.

Hu K, Radhakrishnan P, Patel RV, Mao JJ. 2001. Regional structural and viscoelastic properties of fibrocartilage upon dynamic nanoindentation of the articular condyle. J Struct Biol. 136(1):46–52.

Kuroda S, Tanimoto K, Izawa T, Fujihara S, Koolstra JH, Tanaka E. 2009. Biomechanical and biochemical characteristics of the mandibular condylar cartilage. Osteoarthritis Cartilage. 17(11):1408–1415.

Lei J, Chen S, Jing J, Guo T, Feng J, Ho TV, Chai Y. 2022. Inhibiting hh signaling in gli1(+) osteogenic progenitors alleviates tmjoa. J Dent Res. 101(6):664–674.

Lin DC, Horkay F. 2008. Nanomechanics of polymer gels and biological tissues: A critical review of analytical approaches in the hertzian regime and beyond. Soft Matter. 4(4):669–682.

Ma C, Jing Y, Li H, Wang K, Wang Z, Xu C, Sun X, Kaji D, Han X, Huang A et al. 2021. Scx(lin) cells directly form a subset of chondrocytes in temporomandibular joint that are sharply increased in dmp1-null mice. Bone. 142:115687.

Melou C, Pellen-Mussi P, Jeanne S, Novella A, Tricot-Doleux S, Chauvel-Lebret D. 2022. Osteoarthritis of the temporomandibular joint: A narrative overview. Medicina (Kaunas). 59(1).

Miyawaki S, Tanimoto Y, Inoue M, Sugawara Y, Fujiki T, Takano-Yamamoto T. 2001. Condylar motion in patients with reduced anterior disc displacement. J Dent Res. 80(5):1430–1435.

Murphy MK, Arzi B, Hu JC, Athanasiou KA. 2013. Tensile characterization of porcine temporomandibular joint disc attachments. J Dent Res. 92(8):753–758.

Nishio C, Tanimoto K, Hirose M, Horiuchi S, Kuroda S, Tanne K, Tanaka E. 2009. Stress analysis in the mandibular condyle during prolonged clenching: A theoretical approach with the finite element method. Proc Inst Mech Eng H. 223(6):739–748.

Petitjean N, Canadas P, Royer P, Noel D, Le Floc’h S. 2023. Cartilage biomechanics: From the basic facts to the challenges of tissue engineering. J Biomed Mater Res A. 111(7):1067–1089.

Porter A, Peng Y, Santare MH, Han L, Peloquin JM, Lu XL. 2025. Microstructure and matrix-filled lacunae impact mechanical response of temporomandibular joint cartilage under physiological loading. Osteoarthritis Cartilage. 33(12):1465–1474.

Pueyo Moliner A, Ito K, Zaucke F, Kelly DJ, de Ruijter M, Malda J. 2025. Restoring articular cartilage: Insights from structure, composition and development. Nat Rev Rheumatol. 21(5):291–308.

Roberts RR, Bobzin L, Teng CS, Pal D, Tuzon CT, Schweitzer R, Merrill AE. 2019. Fgf signaling patterns cell fate at the interface between tendon and bone. Development. 146(15).

Ruggiero L, Zimmerman BK, Park M, Han L, Wang L, Burris DL, Lu XL. 2015. Roles of the fibrous superficial zone in the mechanical behavior of tmj condylar cartilage. Ann Biomed Eng. 43(11):2652–2662.

Shibata S, Suda N, Suzuki S, Fukuoka H, Yamashita Y. 2006. An in situ hybridization study of runx2, osterix, and sox9 at the onset of condylar cartilage formation in fetal mouse mandible. J Anat. 208(2):169–177.

Singh M, Detamore MS. 2008. Tensile properties of the mandibular condylar cartilage. J Biomech Eng. 130(1):011009.

Tanaka E, Yamano E, Dalla-Bona DA, Watanabe M, Inubushi T, Shirakura M, Sano R, Takahashi K, van Eijden T, Tanne K. 2006. Dynamic compressive properties of the mandibular condylar cartilage. J Dent Res. 85(6):571–575.

Tang S, Zhang C, Oo WM, Fu K, Risberg MA, Bierma-Zeinstra SM, Neogi T, Atukorala I, Malfait AM, Ding C et al. 2025. Osteoarthritis. Nat Rev Dis Primers. 11(1):10.

Tsouknidas A, Jimenez-Rojo L, Karatsis E, Michailidis N, Mitsiadis TA. 2017. A bio-realistic finite element model to evaluate the effect of masticatory loadings on mouse mandible-related tissues. Front Physiol. 8:273.

Tsutsumi-Arai C, Arai Y, Tran A, Salinas M, Nakai Y, Orikasa S, Ono W, Ono N. 2024. A pthrp gradient drives mandibular condylar chondrogenesis via runx2. J Dent Res. 103(1):91–100.

Tuwatnawanit T, Anthwal N, Tucker AS. 2025a. Activating endogenous condylar stem cells to enhance tmj repair. J Dent Res. 104(13):1443–1452.

Tuwatnawanit T, Wessman W, Belisova D, Sumbalova Koledova Z, Tucker AS, Anthwal N. 2025b. Fsp1/s100a4-expressing stem/progenitor cells are essential for temporomandibular joint growth and homeostasis. J Dent Res. 104(5):551–560.

Uptegrove A, Chen C, Sahagun-Bisson M, Kulkarni AK, Louie KW, Ueharu H, Mishina Y, Omi-Sugihara M. 2025. Influence of bone morphogenetic protein (bmp) signaling and masticatory load on morphological alterations of the mouse mandible during postnatal development. Arch Oral Biol. 169:106096.

Wang J, Dong X, Lei J, Zhang Y, Chen S, He Y. 2024. Beta-catenin orchestrates gli1+ cell fate in condylar development and tmjoa. J Dent Res. 103(12):1291–1301.

Xu M, Zhang X, He Y. 2022. An updated view on temporomandibular joint degeneration: Insights from the cell subsets of mandibular condylar cartilage. Stem Cells Dev. 31(15-16):445–459.

Xu X, Zhang Y, Zhang J, Wang D, Yang H, Liu Q, Zhou P, Wang Y, Yang L, Wang M. 2023. Zonal interdependence in the temporomandibular joint cartilage. FASEB J. 37(4):e22888.

Zhang Y, Feng X, Zheng B, Liu Y. 2024. Regulation and mechanistic insights into tensile strain in mesenchymal stem cell osteogenic differentiation. Bone. 187:117197.

## Appendix References

Muzumdar MD, Tasic B, Miyamichi K, Li L, Luo L. 2007. A global double-fluorescent cre reporter mouse. Genesis. 45(9):593–605.

